# Multimodal Fusion of Circular Functional Data on High-Resolution Neuroretinal Phenotypes

**DOI:** 10.64898/2026.04.02.716157

**Authors:** Saumyadipta Pyne, Hyeongseong Lee, Brian Wainwright, Md. Hasnat Ali, Meghana S. Ray, Sirisha Senthil, S. Rao Jammalamadaka

## Abstract

Progressive optic neuropathies, particularly glaucoma, represent a significant global health challenge, and the need for precise understanding of heterogeneous neurodegenerative phenotypes cannot be overstated. Here, we brought together two complementary sources of unstructured yet clinically relevant information about neuroretinal rim (NRR) thinning, a common clinical marker of such decay. These are based on a new dataset of fundus digital images and a corresponding dataset of optical coherence tomography, both collected from a large clinical cohort of healthy eyes.

First, we represented them using a common data structure that imposed a high-resolution scale of 180 equally spaced and registered measurements on a 360° circular axis. We modeled the NRR measurements of each eye as circular curves and aligned these multimodal curves to obtain a fused NRR curve for each eye.

Unsupervised clustering of these fused curves identified four clusters of eyes with structural heterogeneity, which were also found to have distinctive clinical covariates. Computation of functional derivatives revealed troughs in the curves of each cluster. Using circular statistics, we estimated the directional distributions of these troughs as potentially clinically relevant regions of NRR degeneration.

A comparative study using landmark registration based on functional canonical correlation analysis demonstrated that our curve-alignment-based multimodal fusion is superior. Moreover, it improves the robustness of baseline NRR data obtained from fundus imaging.

## 1 Introduction

Progressive optic neuropathies, particularly glaucoma, represent a significant global health challenge, constituting the second-leading cause of blindness worldwide. With an estimated 80 million individuals affected globally in 2020, and projections indicating this number will exceed 111 million by 2040, the urgency for precise diagnostic methodologies cannot be overstated (Tham, Li, Wong, Quigley, Aung and Cheng, 2014). Glaucoma is characterized by irreversible damage to retinal ganglion cells and the retinal nerve fiber layer (RNFL), progressive neuroretinal rim (NRR) thinning, and excavation of the optic nerve head (ONH). The insidious nature of this disease presents a critical diagnostic challenge: substantial loss of retinal ganglion cells may occur before functional visual field defects become detectable by standard automated perimetry; in fact, by the time visual field defects manifest, 30% of ganglion cell loss may have already occurred (Kerrigan-Baumrind, Quigley, Pease, Kerrigan and Mitchell, 2000). Hence, the stated challenge is best addressed by monitoring and early detection of the dynamic changes in the *normative* neuroretinal phenotypes.

The advent of spectral-domain optical coherence tomography (SD-OCT) has revolutionized ophthalmic diagnostics. SD-OCT offers exceptional glaucoma diagnostic performance with its non-invasive, precise, and reproducible quantitative measurements of critical anatomical structures including the NRR, RNFL, and ONH parameters (Leung, Cheung, Weinreb, Qiu, Liu, Li, Xu, Fan, Huang, Pang et al., 2009). In past studies, we used the OCT technology to generate novel high-resolution circular measurements of NRR and RNFL around the optic disc for characterizing, in unprecedented detail, the spatial distribution of tissue loss, enabling directionally precise analysis of structural changes (Ali, Wainwright, Petersen, Jonnadula, Desai, Rao, Srinivas, Jammalamadaka S. Rao, Senthil and Pyne, 2021; Ali, Ray, Jammalamadaka S. Rao, Senthil, Srinivas and Pyne, 2023). In particular, NRR represents the portion of the ONH that contains viable nerve fibers, and its progressive thinning serves as an early marker of disease progression. Using clinically invisible landmarks such as Bruch’s membrane opening to standardize NRR measurements, OCT provides objective assessments of tissue integrity (Chauhan, O’Leary, AlMobarak, Reis, Yang, Sharpe, Hutchison, Nicolela and Burgoyne, 2013). Based on our analyses, our high-resolution OCT NRR data on a circular scale allowed for the detection of heterogeneous patterns of tissue loss even in normative neuroretinal phenotypes.

Yet, the more cost-effective and popular baseline diagnostic tool worldwide for glaucoma monitoring is fundus photography. Traditional fundus imaging captures the overall morphology of the optic disc and surrounding retinal structures, while advanced imaging modalities enable detailed analysis of structural relationships and the dynamic changes therein. Motivated by the focused directional insights into disease progression owing to our above-mentioned OCT datasets, we generated, in this study, a new dataset with high-resolution measurements based on fundus retinal images for the same cohort of healthy eyes and defined on the same circular scale. we obtained from a given fundus image of an eye, a sample that consists of *L* = 180 equally-spaced non-negative observations defined on the circular domain of [0, 2*π*] radians. Notably, in lower-resolution data summarized for, say, only 4 quadrants or 12 clock-hours on the same circular scale, it may be much harder to detect early focused alterations in degenerative phenotypes. In fact, monitoring such “troughs” as representative of the dynamic process of NRR loss could be even more effective with multimodal data which provides independent evidence to verify consistency across distinct technologies recording the same phenomenon. Thus, we used a shared circular data structure suitable for multimodal fusion of NRR data from OCT and fundus by aligning their complementary information in a form that — as demonstrated below — is more effective for detecting robust neurodegenerative patterns than that of the baseline fundus imaging approach used commonly.

To analyze data in this particular form, we developed a computational platform, CIFU, which represents high-resolution data as smooth circular functions or curves (Ali et al., 2021). Thus, certain tools of functional data analysis (FDA), an established field of statistics in which data are not analyzed as points but as functions (Ramsay and Silverman, 2005), could be utilized effectively in applications involving circular domains, e.g., circadian gene expression analysis (Wainwright, Ghasemi, Ali, Ray, Pyne and Jammalamadaka S. Rao, 2025). The workflow of CIFU begins by modeling each data-point in the form of *L* discrete observations with an optimal number (*p*) of basis functions that results in a continuous curve *X*^*p*^(*t*) defined on a circular scale, i.e., *t* ∈ [0, 2*π*]. Thus, CIFU can normalize a curve to be of unit length while preserving its shape, which enables systematic comparison of length-normalized curves. Importantly, CIFU can be used for outlier detection as well as model-based clustering of the circular curves. Recently, it was also extended with curve registration capabilities (Wainwright et al., 2025).

Special cases of functional data that are based on the temporal domain such as time series, growth curves, etc., commonly encounter the challenge of curve misalignment. To address this, Elastic Shape Analysis (ESA) separates two orthogonal components of functional variability: amplitude and phase (Tucker, Wu and Srivastava, 2013; Harris, Tucker, Li and Shand, 2021). While phase variability could be just a nuisance parameter in certain problems, here it serves as a source of key directional information involving the circular domain. Such variability involves locally observed phenotypic differences in our multimodal data (OCT and fundus NRR curves) and may be representative of the underlying degenerative processes due to neuroretinal thinning. Hence, we addressed the phase variability of these multimodal curves over the common domain by pooling their information in the form of a fused NRR circular curve for each eye. It is computed under a nonparametric form of the Fisher-Rao metric introduced by Srivastava, Wu, Kurtek, Klassen and Marron (2011). Notably, it is a proper distance metric that is invariant to random warping and used commonly in ESA (Srivastava and Klassen, 2016; Harris et al., 2021). The amount of elastic deformation required to fuse two such curves is given by their phase distance, while the residual L ^2^ distance between them post-alignment gives their amplitude distance (Harris et al., 2021). This makes such fusion the most effective in combining pooling phenotypic patterns across those OCT and fundus curves that have low phase distances over their common circular domain.

In the present study, we introduce new high-resolution fundus NRR data, which we used for multimodal fusion with past OCT NRR data (Ali et al., 2021) of similar resolution based on the same cohort of eyes. Our aims are to (a) represent each of these 2 types of NRR data as circular functions or curves, (b) align these multimodal curves based on their phase distances to compute a fused NRR curve for each eye, (c) do unsupervised functional clustering of the fused curves, (d) identify the functional minima or troughs in the curves of each cluster, (e) perform detailed directional analysis of such troughs as regions of NRR decay in eyes based on their circular properties and clinical covariates within each cluster, and (f) conduct a comparative study using an alternative landmark registration approach based on functional Canonical Correlation Analysis, which demonstrated the superiority of multimodal fusion by curve alignment. We note that it also leads to improvement in the robustness of data obtained from the baseline fundus NRR imaging. The next section describes the data and the methods used for different steps of our analysis, followed by a section on results. Finally, we end with a discussion of our present study and future work.

## 2 Data and Methods

### 2.1 NRR data acquisition and preprocessing

As noted in (Ali et al., 2021), participants for this dataset were drawn from two distinct studies: the LVPEI Glaucoma Epidemiology and Molecular Genetic Study (LVPEI-GLEAMS), which is a population-based study, and the Longitudinal Glaucoma Evaluation Study (LOGES), a cross-sectional study. These studies were both conducted by the L.V. Prasad Eye Institute (LVPEI) located in Hyderabad, India. Ethical approval for data protocols was granted by the LVPEI Institutional Ethics Committee for both LVPEI-GLEAMS (LE-08131) and LOGES (LEC 11-252) prior to their implementation. The methodologies were rigorously reviewed and endorsed by the LVPEI ethics and review committee, adhering closely to the principles outlined in the Declaration of Helsinki. All participants provided informed consent prior to their involvement in the research.

While the OCT NRR dataset based on the stated cohort is described in our earlier study (Ali et al., 2021) and reused here for data fusion, the new fundus NRR high-resolution dataset for a common set of *N* = 668 healthy eyes was generated for the present study. Fundus imaging data were acquired using standardized clinical protocols to ensure consistent illumination, focus, and field of view for all eyes. A Topcon non-mydriatic retinal camera, TRC-NW8 (Topcon, Bauer Drive, Oakland, NJ) was used to acquire digital fundus photographs. Subsequently, the NRR thickness information from fundus images was obtained using an image processing workflow (see Supplementary Figure 1) described in the Supplementary Materials.

### 2.2 Circular functional data analysis

CIFU (Ali et al., 2021) represents a data-point as *L* discrete observations made in a wrapped around domain (at points denoted typically by temporal, spatial or directional coordinates) with the help of a circular function or curve, as we have in the present datasets. It begins with the use of *p* basis functions for capturing the functional nature of data. If *ψ*_1_, *ψ*_2_, *· · ·, ψ*_*p*_ are the basis functions with the associated basis expansion coefficients *γ*_*ij*_, where *i* = 1, 2, *· · ·, n* and *j* = 1, 2, *· · ·, p*, then the functional approximation for the *i*^*th*^ curve at point *t, X*_*i*_(*t*), is given by

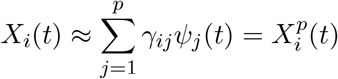

To avoid model overfitting, we determined the smallest number of basis functions (*p*) that recover the input curves sufficiently well, as determined by the fraction of variation explained (FVE) as described in (Ali et al., 2021). Steps for curve normalization and outlier detection are also given in this reference. A subsequent alignment step for fusion of the normalized curves is described below.

Thereafter, CIFU performs curve clustering for unsupervised identification of a pre-specified finite number (*k*) of groups each containing eyes as represented by their aligned curves. Towards this, we used a discriminative functional mixture model (Bouveyron, Côme and Jacques, 2015), which was fit with an Expectation–Maximization (EM) algorithm. The optimal number of clusters (*k*) was determined by multiple well-known model selection criteria such as AIC, BIC and ICL.

### 2.3 NRR curve alignment based fusion

Let Γ be the set of orientation-preserving diffeomorphisms of the unit interval [0, 1] (or [0, 2*π*]), and *f* a real-valued circular function on the same domain. Following (Kurtek, Srivastava and Wu, 2011; Srivastava et al., 2011), the curve alignment method used here represents this function using its square root velocity function (SRVF), where *q* denotes the SRVF of the original function, *q*: [0, 1] *→* ℝ, expressed as

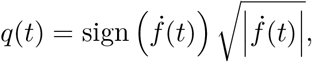

where 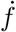 denotes the first derivative of the function *f*. T his a llows alignment of two given curves or functions *f*_1_, *f*_2_ ∈ *ℱ* using the optimal domain warping function *γ*^*^ determined by

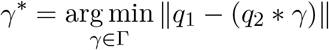

where *q*_1_ and *q*_2_ are the SRVFs of function *f*_1_ and *f*_2_, respectively. In terms of the corresponding SRVFs, *q*_1_, *q*2 *∈* L^2^, their amplitude distance is given by

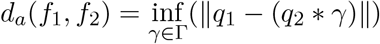

and their phase distance is

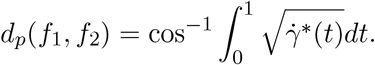

The above definition follows from a nonparametric version of the Fisher–Rao distance using the SRVF framework. We used the R package ‘fdasrvf’ to compute the alignment using a geodesic penalty of the multimodal curves for each eye along with their amplitude and phase distances.

For further details, see (Srivastava and Klassen, 2016).

### 2.4 Density estimation and directional analysis of curve data

Let Θ = {*θ*_1_, *θ*_2_, *· · ·, θ*_*n*_} be a set of circular observations (e.g., functional trough angles), where 0 *≤ θ*_*i*_ *<* 2*π*. The probability density function *f* (*t*) over the angular domain *t* is estimated using an adaptive kernel method to account for varying data density. Instead of Gaussian kernels which are traditionally used for linear data, we employ the von Mises distribution (Jammalamadaka S. Rao and SenGupta, 2001), defined as

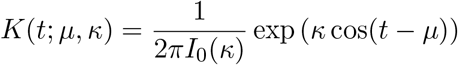

where *µ* is the mean direction, *κ* is the concentration parameter, and *I*_0_(*κ*) is the modified Bessel function of order zero. To capture both sharp peaks and broad distributions, we utilize an adaptive bandwidth strategy. The adaptive estimator 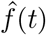 is given by

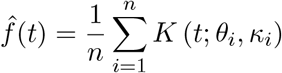

where the local concentration parameter *κ*_*i*_ varies for each observation *θ*_*i*_. The adaptation is controlled by a pilot density estimate 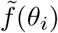 and a global smoothing parameter *h*_0_ (derived via least squares cross-validation). The local bandwidth factor *h*_*i*_ is calculated as

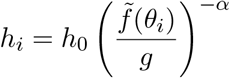

where *g* is the geometric mean of the pilot density values, and *α* ∈ [0, 1] is the sensitivity parameter (typically *α* = 0.5). The local concentration *κ*_*i*_ is then derived inversely from the squared local bandwidth, such that

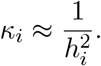

To identify the specific locations of high fusion activity (local modes), we analyze the derivatives of the estimated functional curve 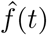. To ensure numerical stability, these derivatives are computed using a B-spline basis expansion. The set of all valid peak locations *P* is defined as

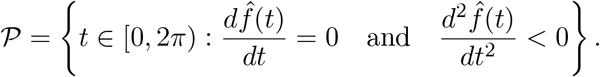

Beyond identifying local peaks, analyzing the variation and centrality of a sample of functional curves requires robust visualization techniques. The functional boxplot (Sun and Genton, 2011) is a natural extension of the classical univariate boxplot to functional data space. Its derivation follows three primary steps: data ordering via depth, identification of descriptive statistics, and outlier detection. To order a sample of *n* curves *X*_1_(*t*), *· · ·, X*_*n*_(*t*) from the center outward, we compute their depths using the Modified Band Depth (MBD). For a given curve *X*(*t*), define the band generated by curves *X*_*i*_ and *X*_*j*_ as *B*_*ij*_(*X*) = {*t* ∈ *ℐ*: *X*(*t*) ∈ [min{*X*_*i*_(*t*), *X*_*j*_(*t*)}, max{*X*_*i*_(*t*), *X*_*j*_(*t*)}]}. Then the sample MBD using bands delimited by *J* = 2 curves is defined as

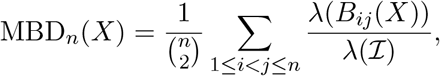

where *λ* denotes the Lebesgue measure and *ℐ* is the angular domain [0, 2*π*]. The sample curves are subsequently ordered according to decreasing depth values, denoted as *X*_[1]_(*t*), *X*_[2]_(*t*), *· · ·, X*_[*n*]_(*t*), where *X*_[1]_(*t*) represents the median or most central curve, and *X*_[*n*]_(*t*) represents the most outlying curve.

Based on this hierarchical ordering, the functional boxplot visualizes three primary descriptive statistics: the median curve, the 50% central region, and the maximum non-outlying envelope. Let *X*_[1]_(*t*), …, *X*_[⌈*n/*2⌉]_(*t*) denote the 50% deepest curves, and define

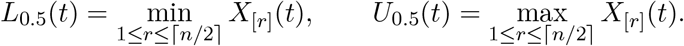

Analogous to the inter-quartile range (IQR), the 50% central region is then

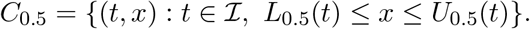

The border of this region forms the “box” of the plot, while the “whiskers” extend from the box to represent the maximum non-outlying envelope of the data. Finally, outliers are detected using an empirical rule analogous to the 1.5 *×* IQR rule. The boundaries, or fences, for outlier detection are calculated by inflating the 50% central region envelope by a factor of 1.5 times its range, defined as

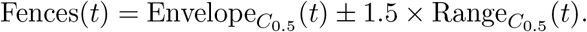

Any curve that falls outside these constructed fences at any point *t* is consequently flagged as a potential outlier.

Finally, to quantify the relationship between the extracted angular measurements, the circular correlation coefficient *ρ*_*cc*_ (Jammalamadaka S. Rao and Ramakrishna Sarma, 1988) *is utilized to measure the association between two angular variables θ* and *ϕ*. It is defined as

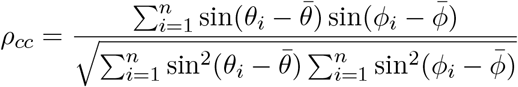

where 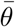 and 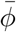 represent the circular mean directions of the respective samples. This correlation coefficient ranges from *−*1 to 1, where a value of *ρ*_*cc*_ *≈* 1 indicates a strong positive circular relationship, and a value of *ρ*_*cc*_ *≈* 0 signifies circular independence.

### 2.5 Comparative Analysis using Landmark Registration and functional CCA

For comparative evaluation of the results of SVRF alignment-based NRR curve fusion, we used another well-known method based on registration of landmark features in functional data (Ramsay and Silverman, 2005). Thus, we used the same multimodal pairs of circular functional NRR data of the non-glaucomatous eyes and computed the corresponding landmark-registered curve using the steps described below. Functional version of canonical correlation analysis (fCCA), a popular method in data fusion, was used for comparaing the results of the wo approaches. For each eye, the OCT-derived NRR curve was treated as the fixed reference curve, while the fundus-derived NRR curve was treated as the source curve to be registered.

Let *i* = 1, …, *n* index eyes, and let *T* = {+91, …, 180} denote the common angular grid. For eye *i*, let

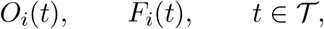

denote the OCT and fundus NRR curves, respectively. Each curve was first area-normalized to remove scale differences:

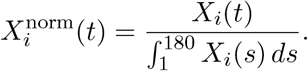

The normalized OCT and fundus curves were then smoothed using a Fourier basis with 15 basis functions over the interval [1, 180]. We denote the resulting smoothed curves by *Õ* _*i*_(*t*) and 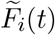.

For each smoothed curve, two peak landmarks were identified. The first landmark was defined as the dominant peak in the first half of the angular domain, excluding the endpoint, and the second landmark was defined as the dominant peak in the second half of the domain, excluding the endpoint. Specifically, for a generic smoothed curve 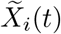,

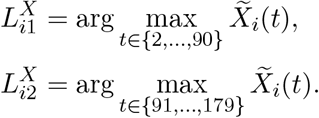

Thus, 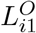 and 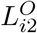 denote the OCT landmarks, while 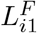 and 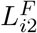 denote the fundus landmarks.

A monotone piecewise-linear warping function was then constructed to align the fundus landmarks to the corresponding OCT landmarks. Define the OCT target landmark vector and fundus source landmark vector as

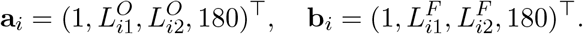

The warping function *h*_*i*_: [1, 180] *→* [1, 180] was defined by

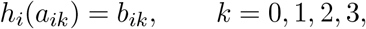

and was linearly interpolated between consecutive landmarks:

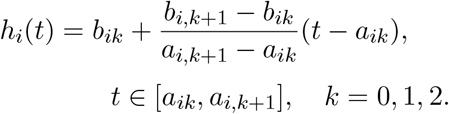

Because the landmarks are ordered within the angular domain, *h*_*i*_ is monotone increasing. The registered fundus curve was obtained by backward warping:

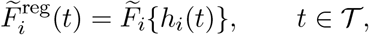

with interpolation used when *h*_*i*_(*t*) did not coincide with an observed grid point. This construction relocates the dominant fundus peaks to the corresponding OCT landmark locations while keeping the OCT curve fixed.

The landmark-registered fused curve was then defined as the equal-weight average of the OCT reference curve and the registered fundus curve:

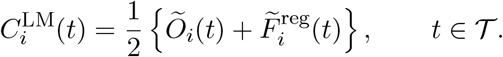

The original fused curve, denoted by 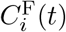, was used as the baseline fusion result for comparison.

To quantify the associations among three sets of curves: (a) the original fundus data modeled as circular functions, (b) fused by curve alignment, and (c) fused by landmark-registration, we used functional CCA (fCCA). Let these sets of curves be respectively denoted by

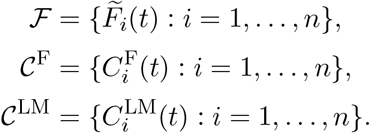

Only the eyes having matched curves across all three sets were included in fCCA. For two functional curve sets *X*(*t*) and *Y* (*t*), fCCA seeks weight functions *α*(*t*) and *β*(*t*) that maximize

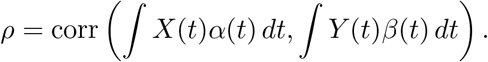

The leading canonical correlations therefore measure the strength of shared subject-level variation between two curve sets. We performed the following pairwise fCCA comparisons:

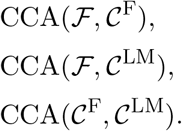

The primary comparison was made between CCA(*ℱ, C*^F^) and CCA(*ℱ, C*^LM^), which evaluates whether the alignment fused curves or the landmark-registered fused curves preserve stronger fundus-detected modes of variation. In addition, we also computed the baseline correlations between the original input curves as well as as the output fused curves. Canonical correlations were summarized using the first canonical correlation, the mean of the first three canonical correlations, and the mean across all reported canonical components.

## 3 Results

We analyzed two high-resolution multimodal datasets: a newly generated fundus-based NRR dataset for this study and a previous one based on OCT, also on NRR, of the same cohort of eyes through a systematic workflow of multiple steps. Each dataset represents an eye as a point comprising *L* = 180 discrete observations made at every 2 degrees of the circular scale. Known landmarks in the eye were used during fundus image preprocessing to match the starting points (0^°^) of both datasets. CIFU workflow was then used to represent the data-points of each type as circular curves, their lengths were normalized, and all outlier curves removed. For each eye, the phase and amplitude distances were computed between its NRR OCT and fundus curves, and results are plotted in Figure 1. The multimodal pair of curves for each eye were aligned using the SVRF methodology and a mean circular function was output as the fused NRR curve (see Supplementary Materials).

Interestingly, whereas there is a unimodal marginal distribution of amplitude distances, that of the phase distances reveal a mixture of points in Figure 1. When fitted with a finite mixture model of 2 components, these points appear as a distinct subset of eyes (shaded green by the posterior probability of component membership) relatively with low phase distances. In particular, most of these points had phase distance less than the median phase distance of 0.447. Thus, we used this value as a threshold, i.e., the points lying to the left of the vertical line displayed in Figure 1 were selected, and only the (*N* = 305) eyes represented by these points were used for the subsequent steps of our analysis.

**Figure 1:**
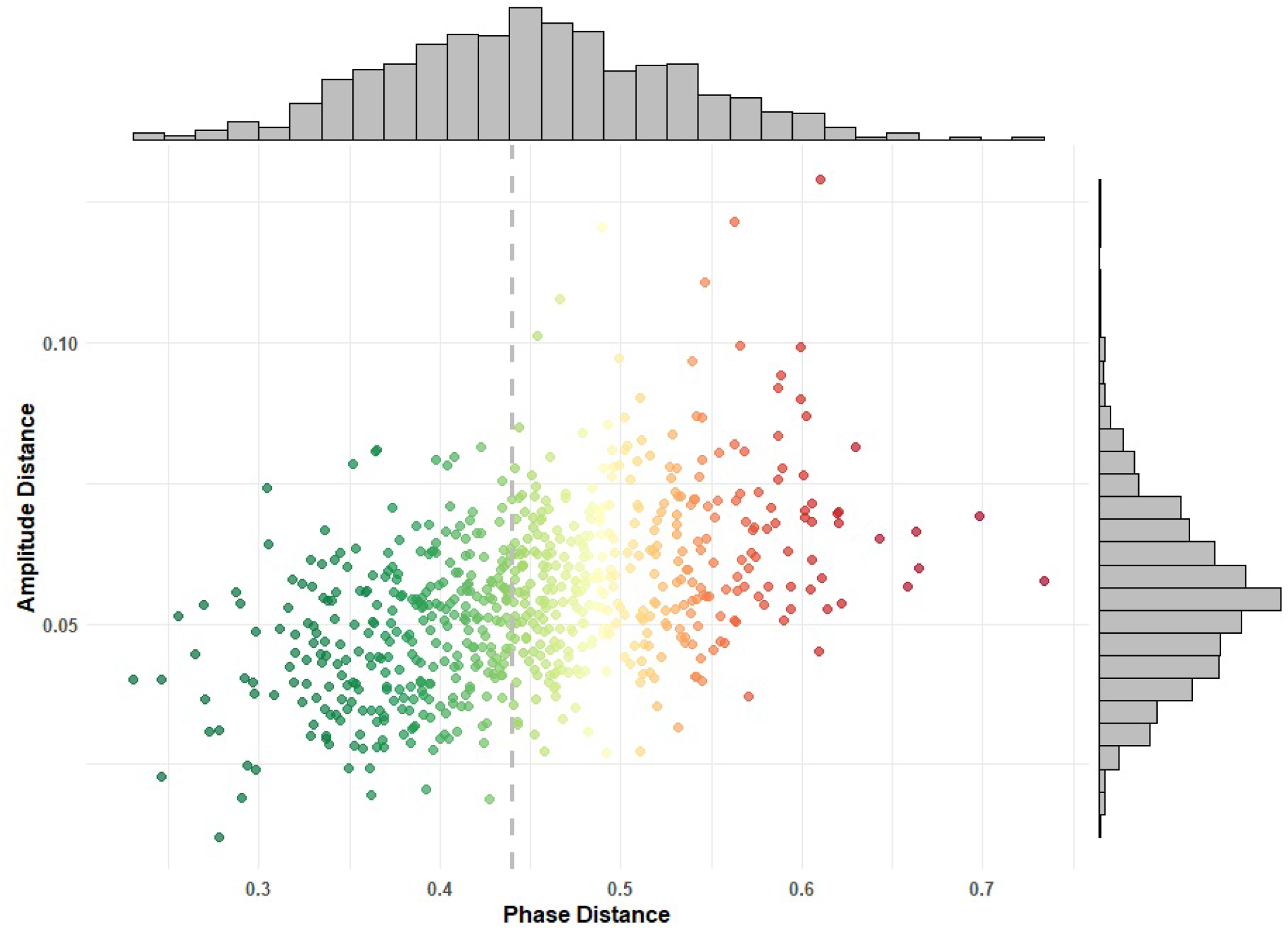
An eye is represented by a point whose x-& y-coordinate respectively denotes the phase and amplitude distances between its NRR OCT and NRR fundus curves. Marginal histograms depict the distributions of each of these distances. The median phase distance is marked as a threshold to select eyes for downstream analyses.

Thereafter, we ran CIFU for unsupervised model-based circular functional data clustering on all the fused curves. Three different commonly-used model-selection criteria (AIC, BIC, and ICL) were computed, and all of these agreed on a final model with 4 clusters of curves, as shown in Supplementary Figure 2. The results of the fused curve clustering are visualized in Figure 2, and the descriptive statistics of each cluster are given in Table 1. The distinction of certain covariates across the clusters such as optic disc area or optic cup volume reveals potential morphologic basis of differential patterns of NRR loss (see Table 1 and Supplementary Figure 3). The mean curve of each cluster is plotted as a black curve that serves as a template for comparative understanding of the overall neurodegenerative patterns. In particular, the cluster means not only allow us to gauge intercluster differences, but also gain insights into intracluster variation, which is also summarized by the trace of the cluster-specific covariance matrix in Table 1.

**Figure 2:**
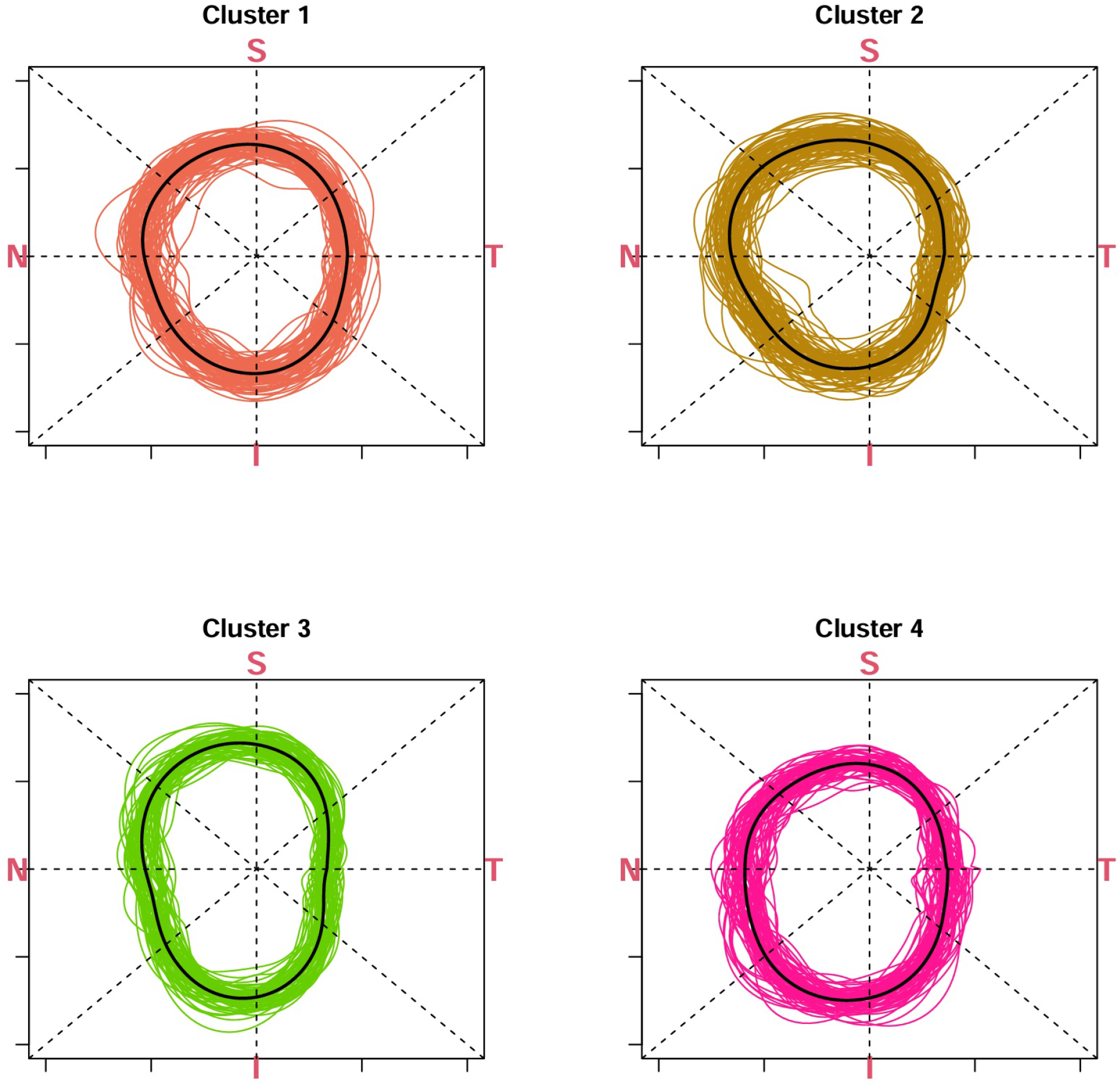
The four clusters of the normalized and fused NRR curves are shown, each in its specific color. For each cluster, its mean NRR curve is shown in black.

**Table 1:** Descriptive Statistics due to CIFU Clustering. The top eleven variables are described as mean *±* standard deviation. (Abbr. BCVA LogMAR: best corrected visual acuity logarithm of the minimum angle of resolution; IOP: intraocular pressure; CCT: central corneal thickness; CD: optic cup-to-disc; *µ*: circular mean; *κ*: circular concentration; OD: oculus dexter; OS: oculus sinister)

| Variable | Cluster 1 | Cluster 2 | Cluster 3 | Cluster 4 |
| --- | --- | --- | --- | --- |
| | $N = 87$ | $N = 84$ | $N = 65$ | $N = 69$ |
| Age (years) | 53.72 ± 10.03 | 54.10 ± 10.09 | 52.28 ± 8.80 | 51.25 ± 8.55 |
| BCVA LogMAR | 0.04 ± 0.08 | 0.07 ± 0.10 | 0.02 ± 0.07 | 0.03 ± 0.08 |
| IOP (mmHg) | 12.22 ± 2.16 | 12.77 ± 2.36 | 12.29 ± 2.48 | 12.32 ± 2.17 |
| CCT (μm) | 523.03 ± 34.64 | 520.70 ± 30.01 | 528.63 ± 30.14 | 521.01 ± 36.43 |
| Axial Length | 22.65 ± 0.86 | 22.50 ± 0.75 | 22.43 ± 0.67 | 22.53 ± 0.79 |
| Rim Area (mm<sup>2</sup>) | 1.34 ± 0.24 | 1.30 ± 0.21 | 1.32 ± 0.20 | 1.37 ± 0.21 |
| Disc Area (mm<sup>2</sup>) | 2.08 ± 0.62 | 2.02 ± 0.52 | 2.26 ± 0.54 | 2.37 ± 0.59 |
| Disc Diameter (mm) | 1.53 ± 0.25 | 1.53 ± 0.21 | 1.56 ± 0.20 | 1.64 ± 0.23 |
| Vertical CD Ratio | 0.46 ± 0.21 | 0.52 ± 0.14 | 0.54 ± 0.12 | 0.57 ± 0.12 |
| Average CD Ratio | 0.52 ± 0.20 | 0.55 ± 0.15 | 0.61 ± 0.11 | 0.61 ± 0.12 |
| Cup Volume (mm<sup>2</sup>) | 0.20 ± 0.20 | 0.24 ± 0.25 | 0.27 ± 0.21 | 0.31 ± 0.22 |
| Sex %(female:male) | 54:46 | 57:43 | 68:32 | 71:29 |
| Eye %(OD:OS) | 52:48 | 27:73 | 26:74 | 52:48 |
| Trace variance | 6.181396e-05 | 7.354637e-05 | 4.768298e-05 | 5.911651e-05 |
| Est. $\mu$ (radians) | 6.17 | 6.13 | 6.13 | 0.04 |
| Est. $\kappa$ | 2.17 | 10.36 | 7.47 | 7.70 |

The circular curve visualization of the clusters allows us to make several interesting observations of various NRR phenotypes both at global and focal scales. Firstly, we find clear evidence for the well-known ISNT rule [34] which specifies that the rim is the thinnest, from the center of ONH, at the temporal (T) region. There is a consistent flatness at the temporal (T) region near 0 degrees, a characteristic shared by the templates of all 4 clusters. Conversely, the “double hump” pattern is evident in the superior (S) and the inferior (I) regions. We note that these features are supported via alignment of more than one data type here. Further, we can observe other interesting shapes and features in these templates that are sometimes shared, e.g., the sudden change in derivative around the nasal (N) region, and at other times, distinctive. The lack of any sharp NRR decay between the nasal (N) and inferior (I) regions in the template of Cluster 4, unlike that in the other clusters, is an example of the latter case. Such distinctions become even more insightful with the inclusion of clinical covariates associated with the eyes in the given clusters, as shown in the Supplementary Figure 3.

Functional box-plots for the fundus and fused curves across each CIFU cluster are displayed in Figure 3. While the fundus distributions exhibit notable outliers, the fused curves consistently provide a smoother, more robust functional representation. It demonstrates that the results of multimodal fusion improves upon the robustness of baseline data obtained from fundus NRR imaging.

**Figure 3:**
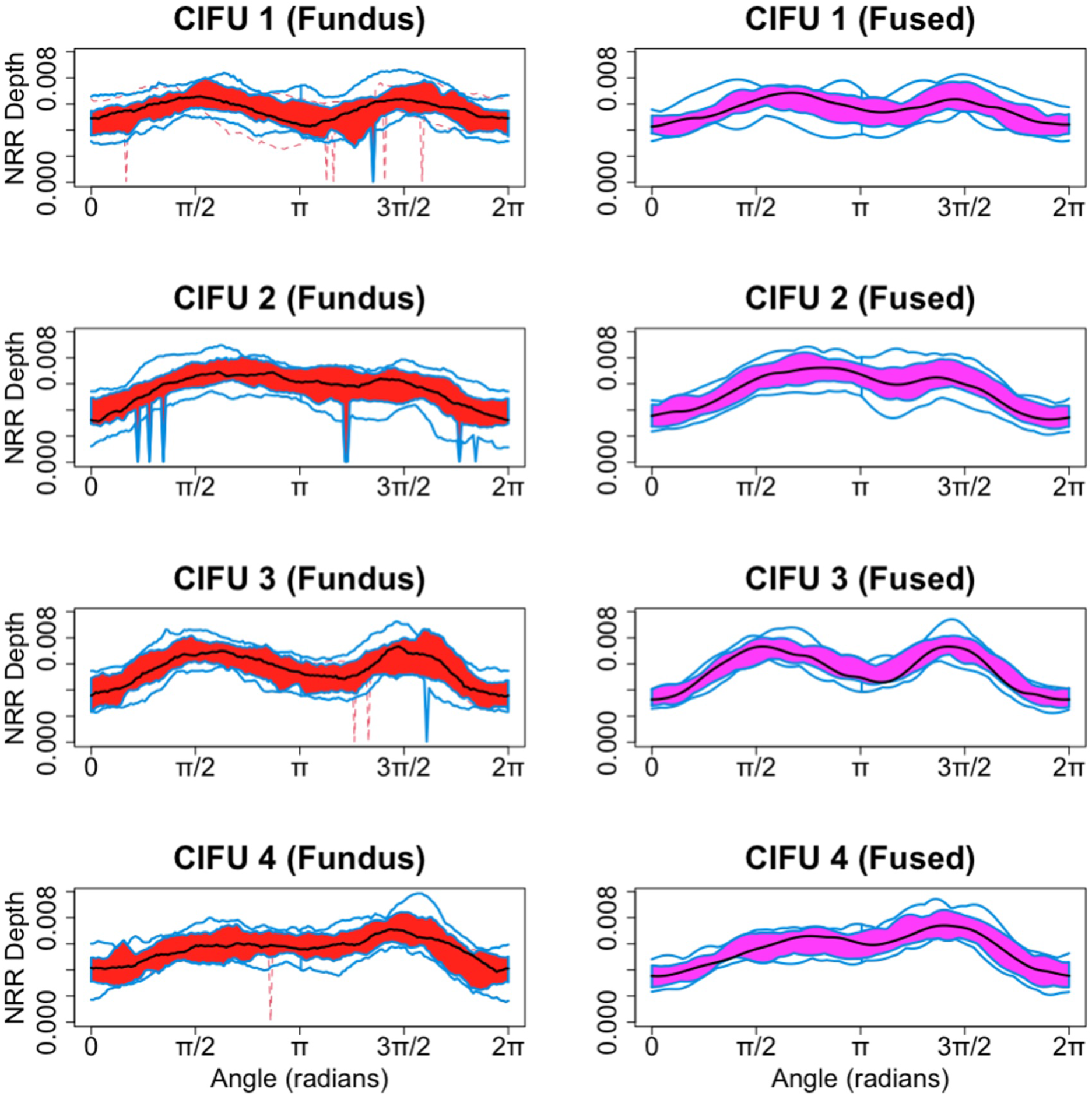
Functional boxplots of fundus (left) and fused NRR curves (right) of each cluster.

To identify the troughs representing regions of NRR loss, we computed the first and second derivatives of the fused curves for each CIFU cluster across the 0 to 2*π* angular domain, as illustrated in Supplementary Figure 4 The tightly bundled trajectories in both the first and second derivative plots confirm the stability and smoothness of the fused representation. Furthermore, the distinct zero-crossings in the first derivative explicitly identify the locations of local maxima and minima, revealing unique, cluster-specific morphological patterns across the angular domain.

The estimated circular kernel densities of the functional troughs for both fundus and fused NRR curves across the four CIFU clusters are presented in Figure 4. Overall, the fused curves demonstrate sharper, more concentrated density peaks near the 0 and 2*π* angular boundaries compared to the wider, more dispersed distributions of the fundus curves. Furthermore, the raw fundus data exhibits noticeable secondary peaks around *π*, most prominently in the CIFU clusters 1 and 3.

**Figure 4:**
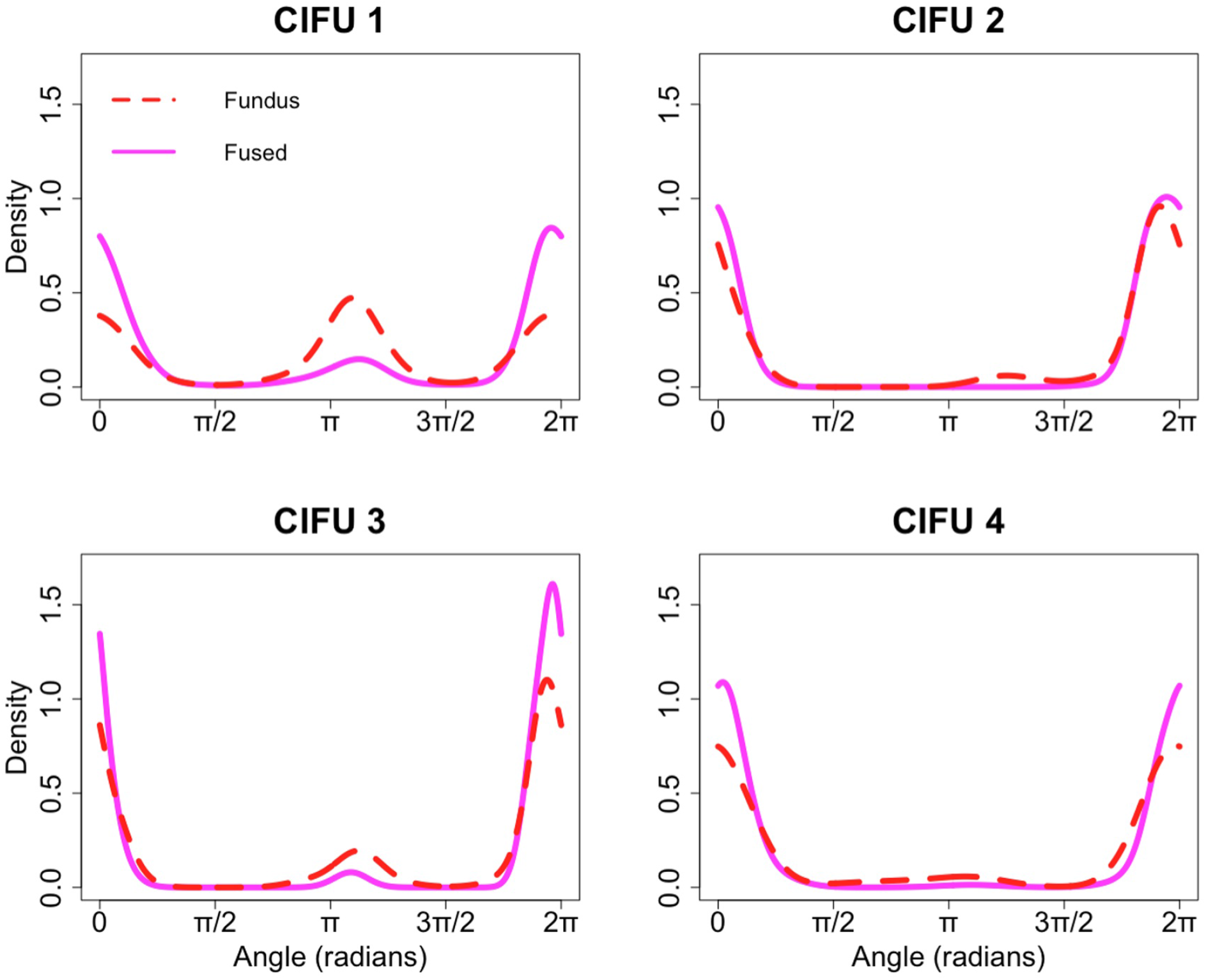
Estimated circular kernel densities of functional troughs in fundus (dashed) and fused (plain) NRR curves of each cluster.

The angular directions of functional troughs (minima) derived from the fundus and fused NRR curves are plotted in Supplementary Figure 5. Due to the periodic nature of the circular data, the dense point clusters located at all four corners of the plot — near 0 and 2*π* on both axes — represent the exact same angular region, demonstrating a strong baseline agreement between the two modalities at the primary structural troughs. We have the circular correlation of 0.3863 (with p-value approximately zero), implying that the functional troughs in the fundus curves are highly associated with the minima in the fused curves.

For comparative analysis, we analyzed the results of multimodal fusion due to curve alignment and landmark registration using functional CCA. The results of landmark registration indicate that the thus fused curves changed the structure of association with the fundus curves rather than uniformly increasing all canonical correlations. As shown in Table 2, landmark-fused curves produce a slightly higher first component of canonical correlation with fundus curves than the alignment fused curves. However, it is the set of alignment fused curves that show larger average canonical correlations over the first three components and, indeed, across all reported components, indicating stronger overall agreement with fundus variation across multiple modes.

**Table 2:** Functional CCA comparison between the NRR curves Fused by the alignment method and those by Landmark registration.

| Criterion | Fused | Landmark-registered | Better method |
| --- | --- | --- | --- |
| First canonical correlation | 0.8656 | 0.8775 | Landmark-registered |
| Mean of first 3 canonical correlations | 0.7935 | 0.7541 | Fused |
| Mean of all reported canonical correlations | 0.3875 | 0.2991 | Fused |

For further insights into the correlation patterns across different canonical components, we see Figure 5. As baselines for comparison, the correlations between the two sets of original pre-fusion NRR curves due to fundus and OCT are displayed (black circles) as well as those between the two sets of fused curves due to alignment and landmark registration (green asterisks). As expected, the green and black lines illustrate the range of correlations that span across the threshold of 0.05 (grey dashed line). Of greater interest are the correlations of the fundus curves with the landmark-registered curves (red triangles) and those with the alignment fused curves (blue squares). Notably, while the former begins with a marginally higher correlation for the first canonical component, it eventually falls below 0.5. In contrast, the latter, i.e., the correlations for alignment fused curves remains above the same threshold. Thus, landmark registration appears to improve the leading fundus-associated mode of variation, while the original fused representation retains broader multi-component similarity to the fundus curves.

**Figure 5:**
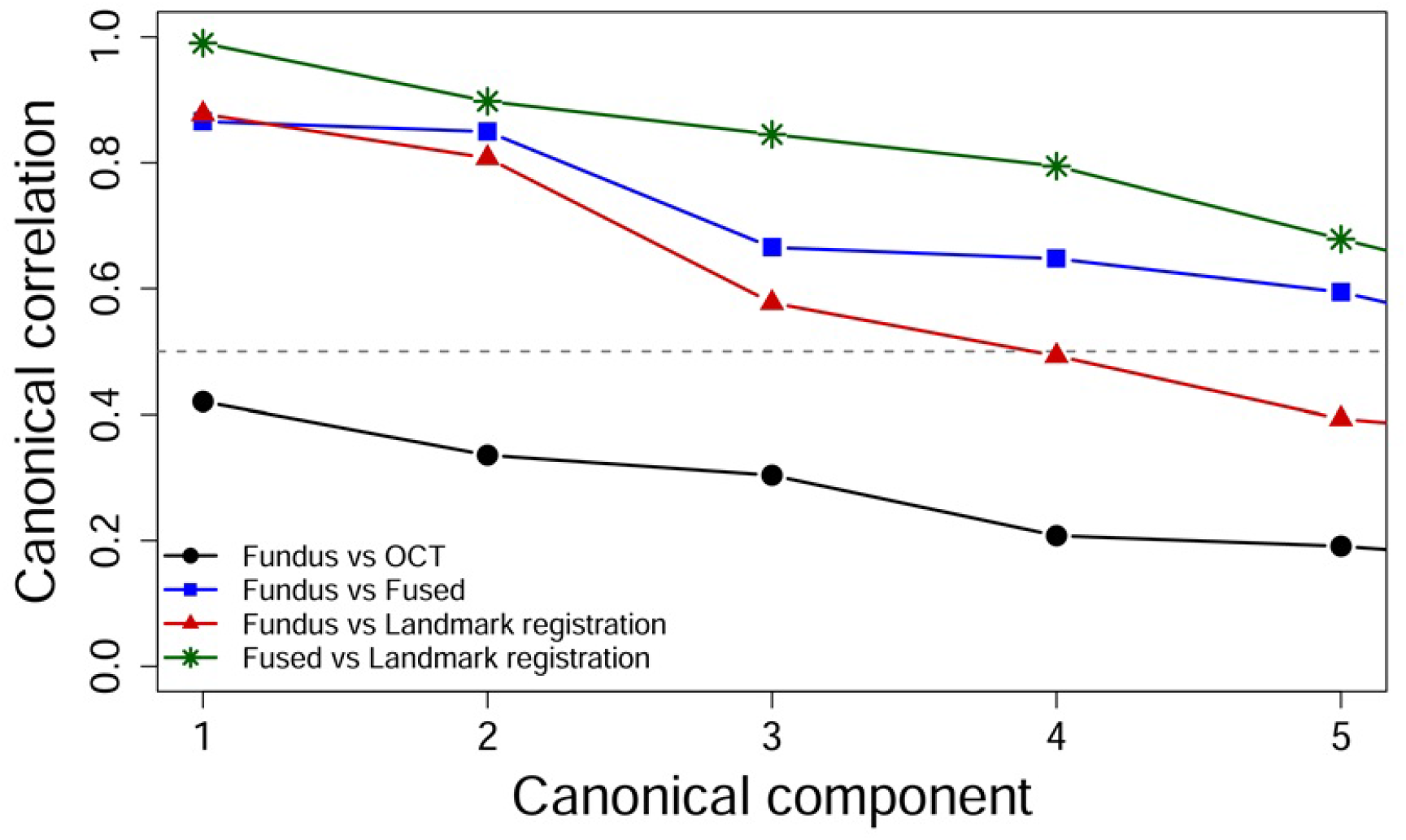
Pairwise functional CCA comparison of NRR curve sets across the first five canonical components. Black circles show Fundus versus OCT, blue squares show Fundus versus alignment Fused curves, red triangles show Fundus versus Landmark-registration fused curves, and green stars show alignment Fused versus Landmark-registration fused curves. The dashed gray horizontal line marks a canonical correlation of 0.5 as a visual reference threshold

## 4 Discussion

In this study, we brought together two different and complementary sources of unstructured but clinically-relevant information — based on eye images and scans — on neuro-retinal phenotypes. First, we represented them using a common data structure that imposed a high-resolution scale of *L* = 180 equally-spaced and registered measurements on the full 360^°^ circular axis. Then, we modeled such NRR data-points of each eye as circular functions or curves. These multimodal curves were aligned to obtain a fused NRR curve for each eye. Unsupervised model-based clustering of these fused curves identified 4 groups of eyes with structural heterogeneity, which were also found to have multiple distinctive clinical covariates. Computation of functional derivatives revealed the crests and troughs in the curves of each cluster. Using a workflow based on circular statistics, we estimated the directional distributions of such troughs as potentially clinically-relevant regions of NRR decay. CCA and its variants have many applications in multimodal data fusion (see review by (Calhoun and Sui, 2016)). Here, we used functional CCA to conduct a comparative analysis, which demonstrated the superiority of multimodal fusion by curve alignment. Finally, using functional boxplot analysis, we demonstrated that the multimodal fusion leads to improvement in the robustness of baseline data obtained from fundus NRR imaging.

Importantly, we presented a rich dataset with a unique combination of three key characteristics: granularity (high-resolution at 180 observations per sample), directionality (covering the entire 360^°^ circular scale), and multimodality (harmonizing digital fundus photographs and OCT scans). Curiously, none of these insightful characteristics of eye images or scans are commonly harnessed in conventional scientific analyses or clinical diagnostics of degenerative neuropathies. We have demonstrated here the utility of such data by first identifying phenotypically heterogeneous clusters, and then estimating within each cluster the directional distributions of troughs, which reveal focal phenotypes of NRR loss. Such focal patterns could be investigated even further with downstream analyses, e.g., with generative or predictive AI models. This could pave the way for future applications of advanced methods in directional and circular statistics (Jammalamadaka S. Rao and SenGupta, 2001) to analysis of optic neuropathies.

We note here that the circular nature of observations has not been entirely ignored in clinical reporting of OCT scans whereby the circle around ONH is divided into 4 quadrants or 12 clock-hours for which the ONH parameters are summarized. Past studies focused on analysis of OCT data summarized over such angular blocks have revealed differences between healthy and glaucoma subjects (Hwang and Kim, 2012). Whereas in low-resolution data, it may be difficult to detect focal alterations using summarized results within the confines of pre-determined inflexible blocks, our continuous curve representation could be analyzed efficiently and rigorously for subtle patterns and their associations with individual-specific variations using functional PCA and functional regression with random effects (Happ, Scheipl, Gabriel and Greven, 2019).

The integration of fundus and OCT data is a useful step that needs to be systematized for clinical applications. Their technological benefits are complementary — while the former is less expensive and more common, the latter is more precise and reproducible. However, it required precise spatial registration to ensure correspondence between the structural features observed in photographs and the quantitative measurements derived from tomographic scans. Advanced image processing algorithms were employed to align coordinate systems and account for individual variations in optic disc size, orientation, and anatomical landmarks, which could be further mechanized (Ramsay and Li, 1998). This registration process harmonized the morphological features visible in fundus images and quantitative parameters measured through OCT scanning and ensured that the output data-points shared the same circular scale with a common origin. Our alignment step rigorously addressed the phase variability in the fused data. This is a painstaking process and a fully automated workflow could be developed to handle the same tasks.

We understand that the present study has certain limitations. Our datasets, despite their desirable properties noted above, offer cross-sectional rather than prospective insights into the dynamic neurodegenerative phenomenon. In our future work, we plan to extend our directional analysis with predictive spatiotemporal models, especially informed by clinical covariates that we already learned here to be distinctive for certain clusters. Our functional representation of data could benefit from integrative models that combine, for instance, time-to-event data on such clinically important outcomes as the onset of glaucoma in individuals (Volkmann, Umlauf and Greven, 2023). The focus on high-resolution multimodal data fusion is a motivator to include further technologies such as OCT-Angiography, used for imaging the microvasculature of the retina, to provide deeper clinical insights in our future work.

In addition to neuroretinal loss with age merely in structural terms, the naturally prevalent biological heterogeneity of ONH phenotypes, especially across different populations, can present significant challenges to clinical decision-making even in the absence of any neuropathy. Our unsupervised identification of both structurally as well as clinically heterogeneous clusters of normative ONH phenotypes can provide useful insights into the diversity of baseline structures that may exist in a given population. Thus, our findings can aid the clinicians in phenotypic classification of eyes, guided by the directional distributions of NRR loss, resulting in potential reduction of subjectivity and delay in the diagnosis of degenerative neuropathies.

## Notes

### Competing Interest Statement

The authors have declared no competing interest.

### Summary of Updates

Revision contains new results and changes in discussion section.

